# Inferring entire spiking activity from local field potentials with deep learning

**DOI:** 10.1101/2020.05.02.074104

**Authors:** Nur Ahmadi, Timothy G. Constandinou, Christos-Savvas Bouganis

## Abstract

Extracellular recordings are typically analysed by separating them into two distinct signals: local field potentials (LFPs) and spikes. Understanding the relationship between these two signals is essential for gaining deeper insight into neuronal coding and information processing in the brain and is also relevant to brain-machine interface (BMI) research. Previous studies have shown that spikes, in the form of single-unit activity (SUA) or multiunit activity (MUA), can be inferred solely from LFPs with moderately good accuracy. These spiking activities that are typically extracted via threshold-based technique may not be reliable when the recordings exhibit a low signal-to-noise ratio (SNR). Another spiking activity in the form of a continuous signal, referred to as entire spiking activity (ESA), can be extracted by a threshold-less, fast, and automated technique and has led to better performance in several tasks. However, its relationship with the LFPs has not been investigated. In this study, we aim to address this issue by employing a deep learning method to infer ESA from LFPs intracortically recorded from the motor cortex area of two monkeys performing different tasks. Results from long-term recording sessions and across different tasks revealed that the inference accuracy of ESA yielded consistently and significantly higher accuracy than that of SUA and MUA. In addition, local motor potential (LMP) was found to be the most highly predictive feature compared to other LFP features. The overall results indicate that LFPs contain substantial information about the spikes, particularly ESA, which could be useful for the development of LFP-based BMIs. The results also suggest the potential use of ESA as an alternative neuronal population activity measure for analysing neural responses to stimuli or behavioural tasks.

## Introduction

Extracellular recordings are one of the most widely used electrophysiological techniques and have been extensively used for basic neuroscience research (e.g. understanding neural coding and information processing) and clinical applications (e.g. brain-machine interface). The advent and advancement of microelectrode array technology have made it possible to simultaneously record neural signals from a large group of neurons^1–3^. The recorded raw neural signals are composed of two main components: local field potential (LFP) and spikes. The LFPs are typically obtained by low pass filtering the raw neural signals (below ∼300 Hz) and are thought to mainly reflect summed synaptic activity from a local neuronal population (within a radius of at least a few hundred micrometres) around the recording electrode^4–7^. On the other hand, the spikes are extracted by high-pass filtering the same raw neural signals (usually above ∼300 Hz) and subsequent spike processings. Based on these subsequent processings, spikes can be categorised into three types of signals, namely single-unit activity (SUA), multiunit activity (MUA) and entire spiking activity (ESA).

SUA is defined as the timing of spikes (i.e. action potentials) fired by individual neuron and is extracted via threshold crossing followed by unit classification known as spike sorting. MUA, which is in some literature called multiunit spike (MSP), refers to all the detected spikes (without spike sorting) and represents the aggregate spikes from an ensemble of neurons within a radius of ∼140-300 *µ*m in the vicinity of the electrode tip^8–11^. Extracting both SUA and MUA rely on setting the threshold value (manually or automatically) which could be problematic when the recordings exhibit a low SNR or high variation over time. This circumstance is often encountered in chronic recordings where the amplitude of spikes decreases due to tissue responses and/or micromotion of the electrodes^12^. Moreover, threshold-based technique may result in a biased estimate of spiking activity in favour of large neurons (pyramidal neurons), hence leaving the spiking activity of small neuron undetected^13, 14^.

Unlike both SUA and MUA which are represented by a sequence of binary signals, ESA is represented by a continuous signal and reflects an instantaneous measure of the number and size of spikes from a population of neurons around the recording electrode^15, 16^. ESA is obtained through full-wave rectification (i.e. taking the absolute value) followed by low-pass filtering^14^. The term ESA is relatively new^17^ but its underlying principle has existed and been used for several decades^15, 18^. Despite being less popular and less frequently used signal than its counterparts, ESA offers appealing advantages. Its threshold-less and automated processing provides a more reliable and less biased estimate of population spiking activity because it is less sensitive to random high-frequency noise and takes into account the spike contribution from small neurons. A number of studies have demonstrated that ESA can achieve better accuracy and reliability in measuring evoked responses (e.g. receptive field) in the visual cortex of monkeys while receiving various visual stimuli^13, 14, 19–21^.

Understanding the relationship between the LFP and spikes plays a critical role in addressing many issues in neuroscience research, such as neuronal functional organisation and connectivity, neuronal communication between different brain areas, cell assembly formation, neural coding and information processing^22–25^. In addition, it is also relevant for brain-machine interface (BMI), for example, measuring behavioural task-related information within different neural signals^26, 27^ and extracting indirect spiking information features from LFPs in biofeedback based BMI^28^. As LFPs are thought to represent mainly the input to local neuronal networks, while spiking activity represents the output from local neuronal networks^29^, it is conceivable to relate the two signals based on system identification-based inference approach. Previous studies have shown that SUA^28, 30, 31^ and MUA^26, 32^ can be inferred solely from LFPs with moderately good accuracy. So far, however, there has been no study that investigates the relationship between the LFPs and ESA.

In light of this, our present study aims to address the above-mentioned issue by using extracellular recordings from motor cortex area of two macaque monkeys while performing two different tasks: point-to-point task and instructed delay reach-to-grasp task. We assess quantitatively how well ESA can be inferred from LFPs with deep learning, specifically long short-term memory (LSTM) network. We also analyse which features within LFPs are highly predictive of ESA. Furthermore, we compare the inference accuracy of ESA with that of SUA and MUA. Finally, we examine the LFP channel importance/contribution and the effect of the number of LFP channels on the inference accuracy.

## Results

### LFP feature informativeness for ESA inference

We evaluated the informativeness of different LFP features for the inference of ESA using extracellular recordings from a subject while performing a point-to-point task (see Dataset I in Methods). We trained and optimised an LSTM model for each LFP feature separately and then quantified how well this model and LFP feature predict ESA. Figure 2a shows boxplots comparing the inference performance of six LFP features measured in average Pearson’s correlation coefficient (CC). From 910 cases (10 blocks × 91 ESA channels), LMP was found to yield the highest accuracy with CC = 0.683 ± 0.008 (mean ± standard error of the mean (SEM)), while power spectra in the alpha band (shortly referred to as alpha) yielded the lowest accuracy with CC = 0.136 ±0.002. The order of LFP feature from highest to lowest accuracy was LMP > gamma > delta > beta > theta > alpha. The statistical tests performed between a pair of LFP features from all possible combinations revealed that there were statistically significant differences (*** *p* < 0.001) between LFP features (as shown in Figure 2b). Relative to other features, LMP yielded on average 1.43 to 5.15 times better CC as illustrated in Figure 2c. Due to the highest predictive power, LMP was thus selected as the feature for subsequent processing and analysis.

**Figure 1.**
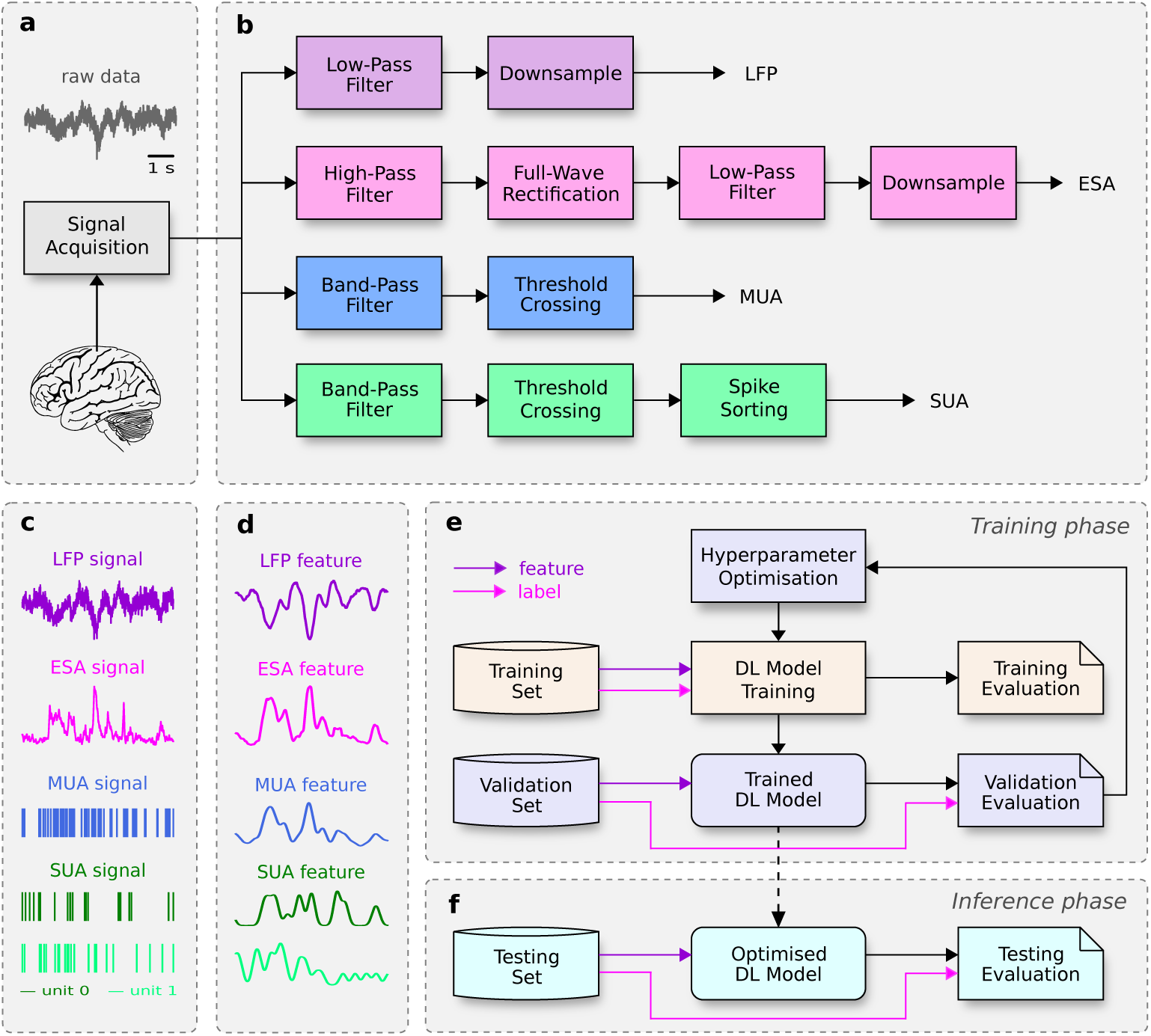
Schematic illustration of signal processing and deep learning (DL) based inference steps. (**a**) Raw neural signal acquisition from the motor cortex area of monkey with a 96-channel intracortical Utah array. (**b**) Signal processing steps for different types of neural signals (LFP, ESA, MUA, and SUA). (**c**) LFP, ESA, MUA, and SUA signals obtained from the processing steps. (**d**) Extracted features from LFP, ESA, MUA, and SUA signals. (**e**) DL model training and optimisation phase. (**f**) DL model inference phase.

**Figure 2.**
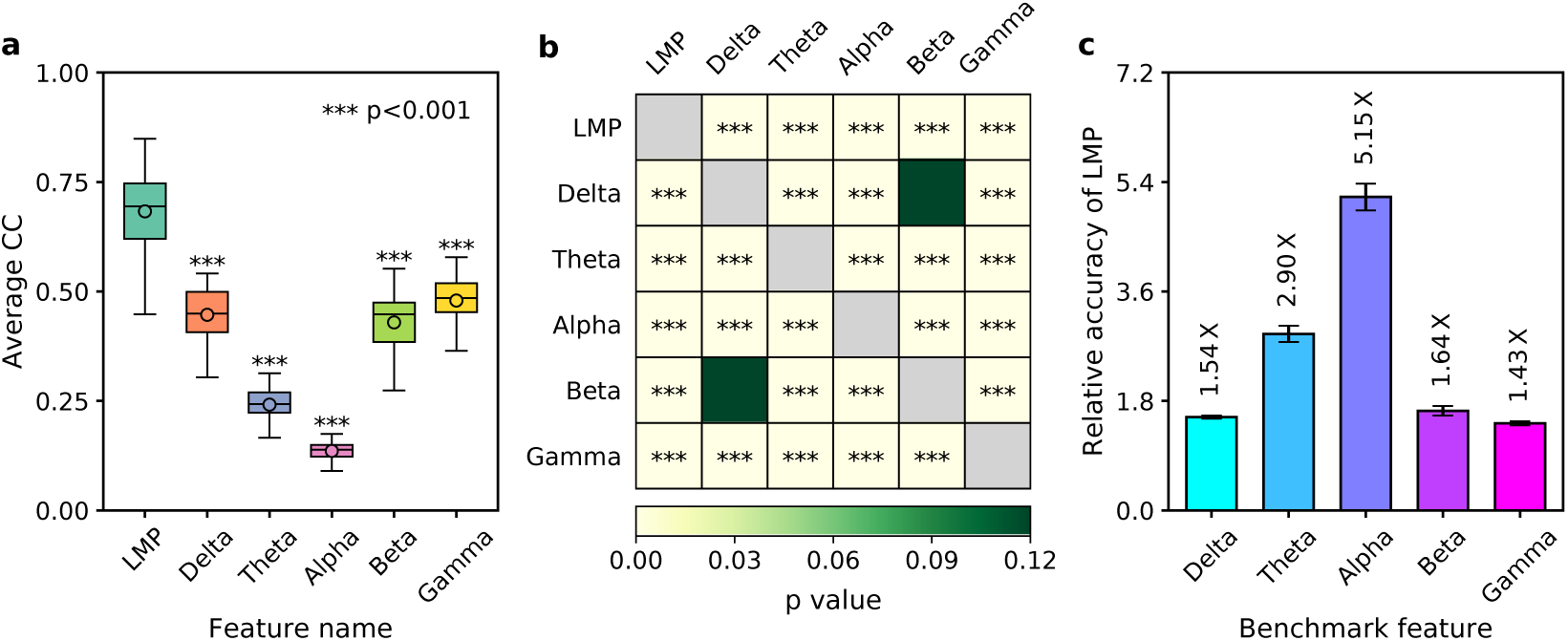
Performance comparison of ESA inference across different LFP features (data from session I20160627_01). (**a**) Boxplot comparison of average CC across LFP features. Asterisks indicate LFP features whose inference performance differed significantly from that of LMP (*** *p*<0.001). (**b**) Heatmap of statistical significance matrix among LFP features. Grey boxes in the main diagonal does not have *p*-value (excluded from the statistical test). (**c**) Relative performance of LMP with respect to other LFP features. Larger value indicates better relative performance. Black error bars denote 95% confidence intervals.

### Comparison of inference performance: SUA vs MUA vs ESA

Using LMP feature from 96 LFP channels in the first recording session of dataset I (I20160627_01), we then performed inference of three different spiking activities: SUA, MUA, and ESA. In this recording session, from a total of 96 channels, we only inferred 83 and 91 channels for SUA and MUA, respectively; other channels were excluded due to very few number of detected spikes (≤0.5 Hz). For the case of MUA, we inferred all the 96 channels but for the performance comparison we only took into account the inference from the same channels as MUA. The inference performance (measured in CC) of each channel of SUA, MUA, and ESA are plotted into heatmap grid spatially corresponding to Utah array configuration as shown in Figure 3a-c, respectively. Numbers within the red boxes in Figure 3a-b indicates the excluded SUA and MUA channels. From 83 SUA channels, we obtained an average CC of 0.428 ± 0.017 (mean ± SEM) with minimum CC of 0.126 and maximum CC of 0.786. From 91 MUA channels, we obtained higher inference accuracy (CC = 0.619 ± 0.014 with CC_*min*_ = 0.071 and CC_*max*_ = 0.858). The highest inference accuracy was obtained from ESA inference with an average CC of 0.685 ± 0.008 across 91 channels with CC_*max*_ = 0.454 and CC_*max*_ = 0.846. The order of spiking activities from highest to lowest inference accuracy was ESA > MUA > SUA.

**Figure 3.**
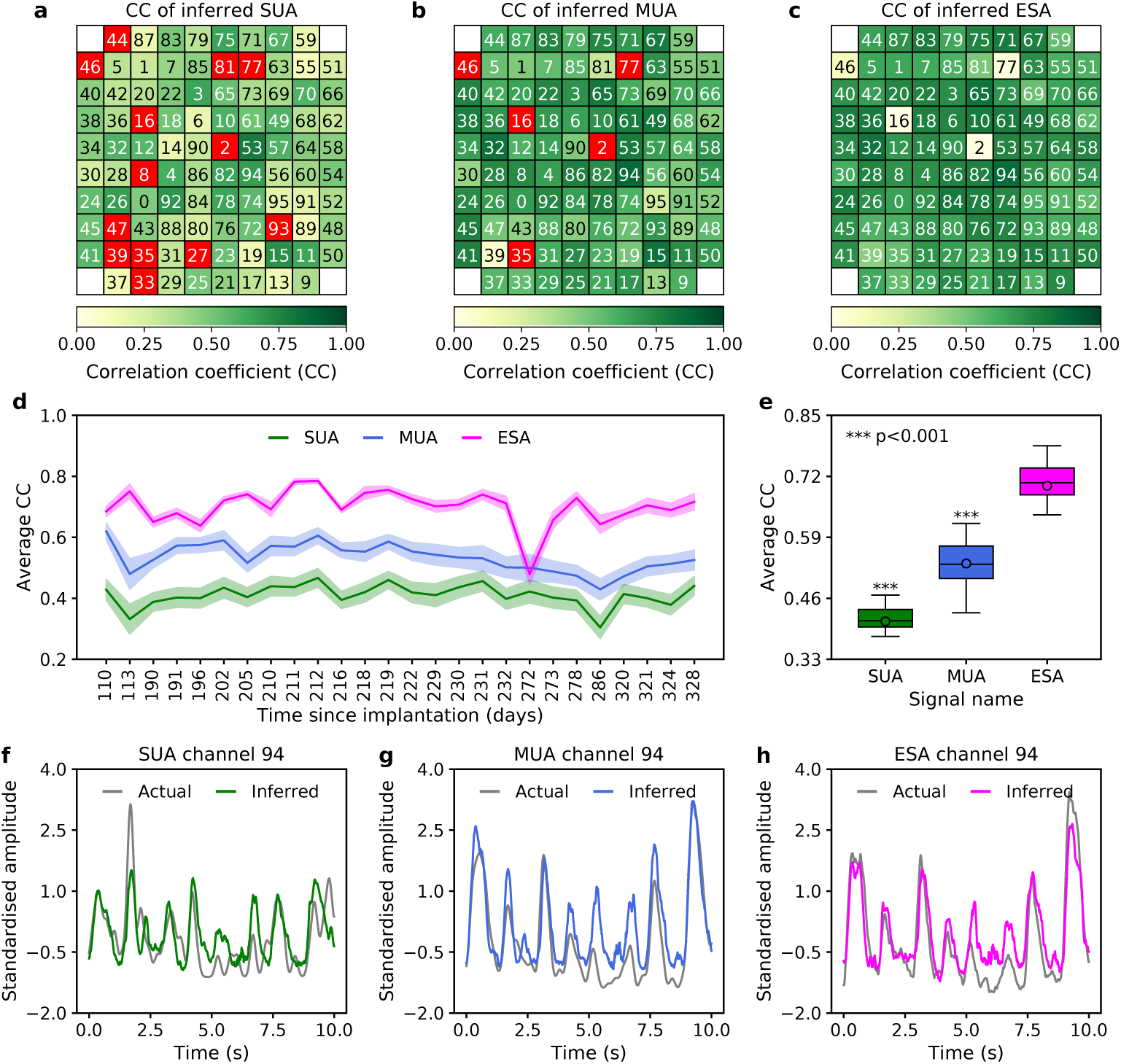
Comparison of inference accuracy (CC) among different types of spiking activities (SUA, MUA, and ESA). (**a**-**c**) Heatmap of CC from 96-channel of SUA, MUA, and ESA, respectively mapped onto a 10 × 10 grid spatially corresponding to Utah electrode array configuration. Black/white numbers inside the grids denote each of the 96-channel numbers. Grey boxes on the grid corners represent unused (unconnected) electrodes. Red boxes indicate channels that were excluded from the inference due to too few number of recorded spikes (≤ 0.5 Hz). Data are taken from session I20160627_01. (**d**) Comparison of average CC among SUA, MUA, and ESA over 26 sessions. (**e**) Boxplot comparison of average CC across sessions among SUA, MUA, and ESA. Asterisks indicate the statistical significance of other spiking activities being compared to ESA (*** *p* < 0.001). (**f**-**h**) A snippet example of actual and inferred SUA, MUA, and ESA, respectively (taken from channel 94).

To investigate whether this trend was consistently observed across different recording sessions, we performed the inference using long-term neural recordings with a total of 26 sessions spanning more than 7 months between the first and last sessions. Across the entire recording sessions, the number of SUA channels being included for analyses varied from 71 to 89 with an average of 80.62 ± 5.36 (mean ± standard deviation (SD)); for both MUA and ESA, the number of included channels were from 77 to 95 channels with an average of 87.08 ± 4.44 (mean ± SD). As can be seen in Figure 3d, the inference accuracy of ESA was consistently better than that of SUA and MUA. Compared to SUA, the inference accuracy of ESA was higher in 26 of 26 sessions (100%), whereas compared to MUA, the inference accuracy of ESA was higher in 25 of 26 sessions (96.15%). The average accuracy (mean ± SEM) of SUA, MUA, and ESA were 0.411 ± 0.007, 0.534 ± 0.009, and 0.700 ± 0.012, respectively. Figure 3e displays boxplot of ESA inference that was statistically significant different than that of SUA and MUA (*** *p* < 0.001). The relative inference accuracy of ESA was 1.71 and 1.32 times higher with respect to SUA and MUA, respectively. These results indicate that LFPs contain substantial information about ESA. Examples of actual and inferred SUA, MUA, and ESA are illustrated in Figure 3f-h.

CC is a translation and scale invariant metric that only assesses the shape similarity between actual and inferred signals. In the presence of a large but constant inference error, the CC value can be high (implying falsely good accuracy). Therefore, as an additional metric for the inference performance, we used root mean square error (RMSE) that represents an average magnitude of inference error between actual and inferred signals. We found the same trend of the LFP feature informativeness as of CC. LMP yielded the highest accuracy (i.e. smallest RMSE) with RMSE = 0.728 ± 0.007 (mean ± SEM), whereas alpha yielded the lowest accuracy (i.e. largest RMSE) with RMSE = 0.987 ± 0.002. The LFP feature informativeness in descending order revealed the same pattern as of CC: LMP > gamma > delta > beta > theta > alpha (see Figure 4a). There were statistically significant differences (*** *p* < 0.001) among combinations of LFP feature pairs as illustrated in a heatmap of *p*-values shown in Figure 4b). In comparison to other features, LMP yielded on average 1.20 to 1.37 times better accuracy (Figure 4c).

**Figure 4.**
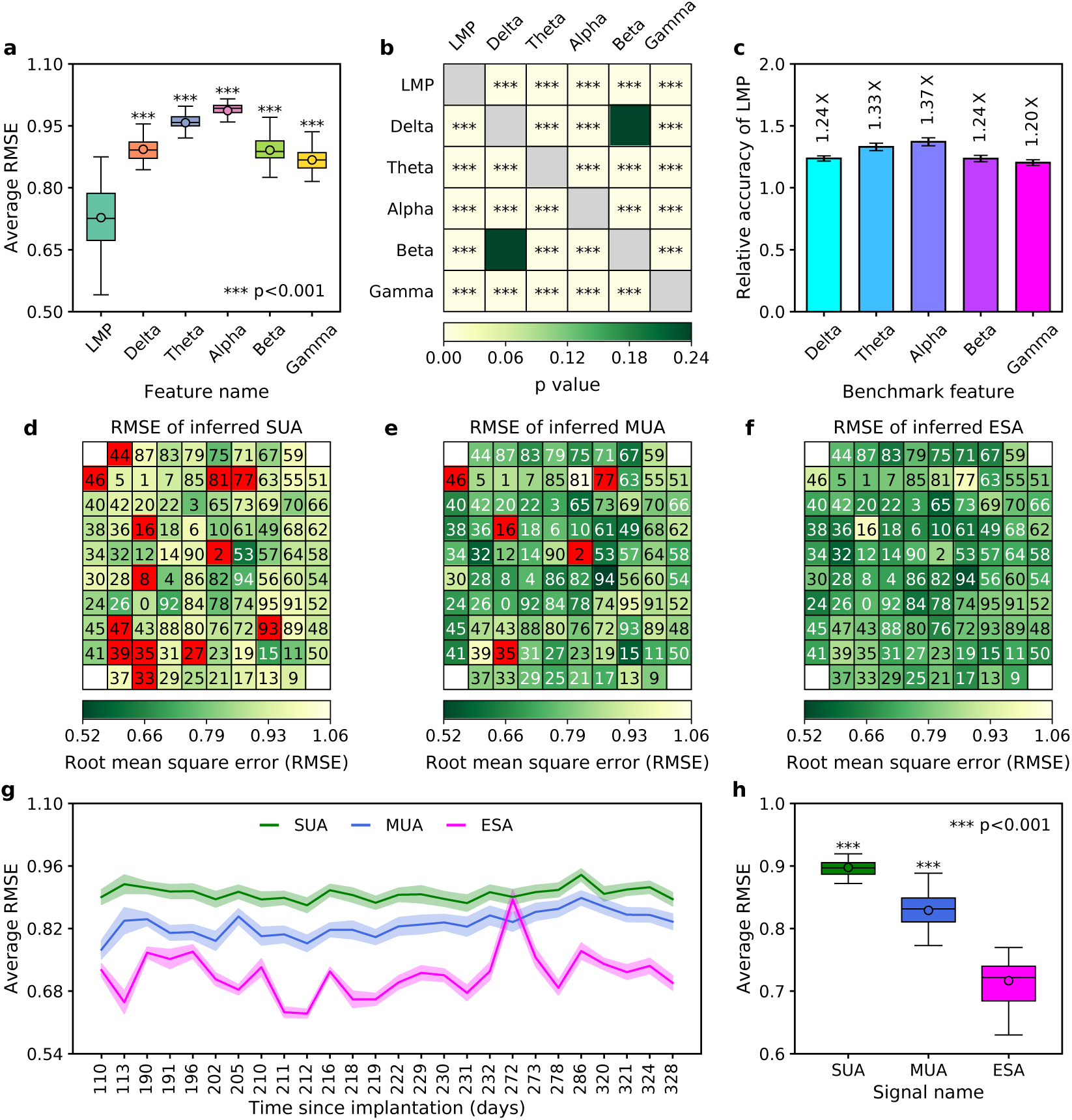
Comparison of inference accuracy measured with RMSE metric. (**a**) Boxplot comparison of ESA inference across LFP features. Asterisks indicate LFP features whose inference performance differed significantly from that of LMP (*** *p*<0.001). (**b**) Heatmap of statistical significance matrix among LFP features. Grey boxes in the main diagonal does not have *p*-value (excluded from statistical test calculation). (**c**) Relative inference accuracy of LMP with respect to other LFP features. Larger value indicates better relative performance. Black error bars denote 95% confidence intervals. (**d**-**f**) Heatmap of RMSE of 96-channel SUA, MUA, and ESA, respectively mapped onto a 10 × 10 grid spatially corresponding to Utah electrode array configuration. Black/white numbers inside the grids denote each of the 96-channel numbers. White boxes on the grid corners represent unused (unconnected) electrodes. Red boxes indicate channels that were excluded from the inference due to too few number of recorded spikes (≤ 0.5 Hz). Data are taken from session I20160627_01. (**g**) Comparison of average RMSE among SUA, MUA, and ESA over 26 sessions. (**h**) Boxplot comparison of average RMSE across sessions among SUA, MUA, and ESA. Asterisks indicate the statistical significance of other spiking activities being compared to ESA (*** *p*<0.001).

Identical to that of CC metric, ESA inference accuracy (average RMSE 0.738 ± 0.008) was better than that of SUA (0.890 ± 0.008) and MUA (0.773 ± 0.010). Figure 4d-f display heatmaps of inference accuracy (RMSE) from each channel of SUA, MUA, and ESA, respectively. Greener grid indicates better accuracy (smaller RMSE) whereas yellower grid indicates worse accuracy (larger RMSE). The order of spiking activities from highest to lowest inference accuracy was ESA > MUA > SUA. As evident from Figure 4g, the same trend was also observed across long-term recording sessions where the inference accuracy of ESA consistently outperformed its counterparts. The average accuracy (mean ± SEM) of SUA, MUA, and ESA were 0.898 ± 0.003, 0.829 ± 0.005, and 0.717 ± 0.010, respectively. Figure 4h displays boxplot of ESA inference showing its statistically significant difference from that of SUA and MUA (*** *p* < 0.001). The inference accuracy of ESA was 1.26 and times better relative to SUA and MUA, respectively.

### LFP channel importance/contribution

Next, we assessed the importance or contribution of LFP channels for the inference of spiking activity at particular channels. The LFP channel importance was measured and ranked using permutation feature importance as described in Methods section. Figure 5a-c show an example of LFP channel importance scores mapped into 10-by-10 heatmap grids for the inference of channel 94 of SUA, MUA, and ESA, respectively. The channel of spiking activity being inferred (channel 94) is displayed in bold yellow text. The LFP channel with highest importance score for the inference of SUA, MUA, and ESA was channel 37, 24, and 94, respectively. From the analysis across different inferred channel numbers, we can draw two insights. First, the LFP from the same channel as the inferred spiking activity on average had high importance score (for example, see Figure 5a-b) than other channels but did not necessarily have the highest importance score (Figure 5a). This indicates that LFP from the same channel contains rich information about spiking activity. Second, for the same channel number, the LFP importance score for ESA inference was higher than that for SUA and MUA. This may indicate that ESA contains additional information that is not available in SUA and MUA.

**Figure 5.**
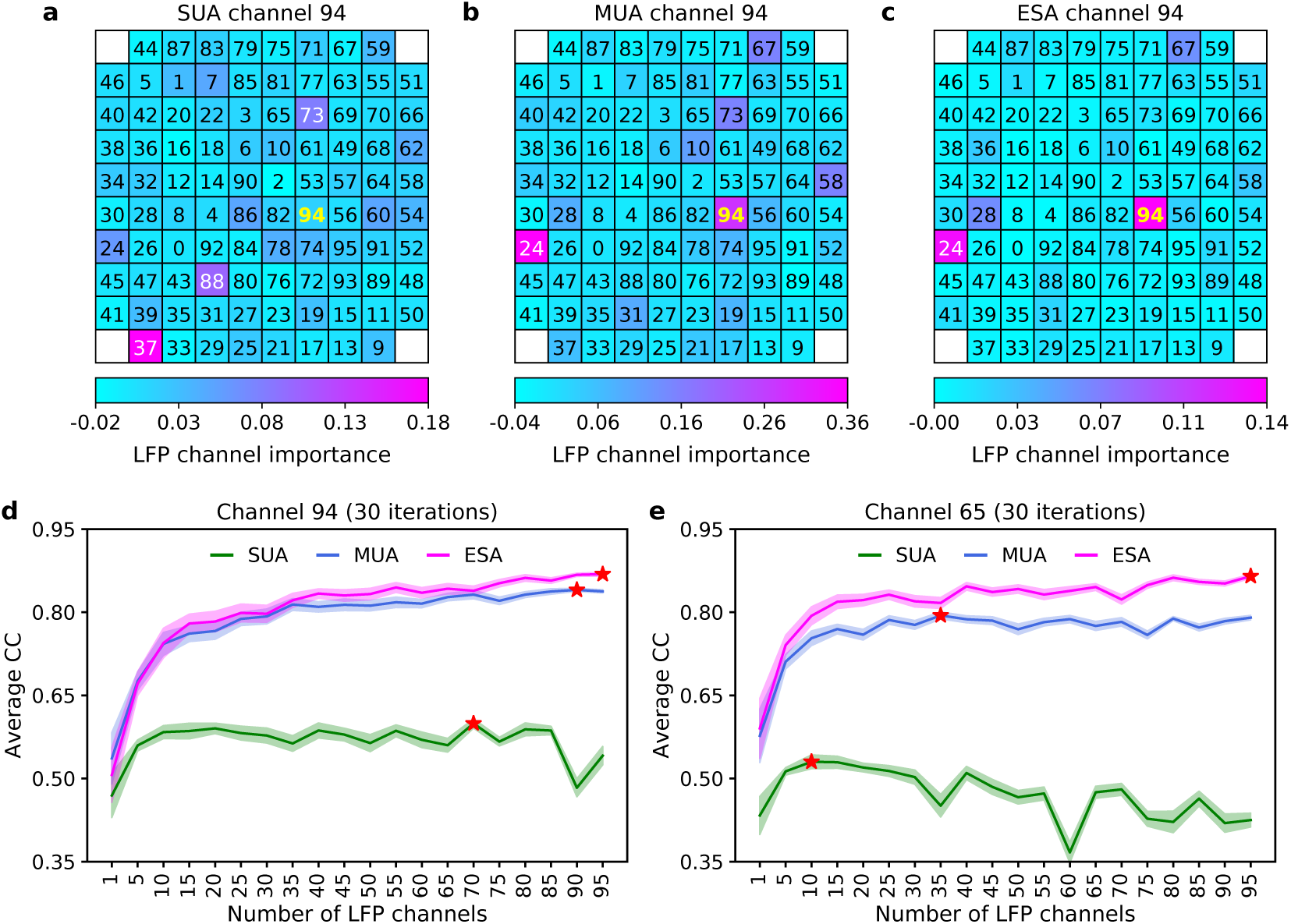
The importance of each LFP channel and the effect of the number of LFP channels to inference accuracy. Data are taken from session I20160627_01. (**a**-**c**) Heatmap of LFP channel importance score for the inference of SUA, MUA, and ESA in channel 94, respectively mapped onto a 10 ×10 grid spatially corresponding to Utah electrode array configuration. Black/white numbers inside the grids denote each of the 96 LFP channel numbers. Bold yellow number (i.e. 94) indicates the channel of spiking activities being inferred. White boxes on the grid corners represent unused (unconnected) electrodes. The larger the channel importance score, the more important is the channel for the inference. (**d**-**e**) The effect of increasing number of LFP channels for the inference accuracy (average CC) of spiking activities in channel 94 and 64, respectively. The shaded areas represent 95% confidence intervals (*N* = 30). The red star markers indicate the number of LFP channels that corresponds to highest average CC score for each spiking activity.

### Effect of number of LFP channels on inference accuracy

Figure 5d-e illustrates the effect of using different number of LFP channels on the inference accuracy. We found that increasing the number of LFP channels to a certain number dramatically improved the inference accuracy. However, after that point, the inference accuracy only improved marginally or even decreased (as evident in the case of SUA from Figure 5d-e). Analysis across different inferred spiking activity channels revealed that the inference accuracy of SUA reached a plateau in significantly less number of LFP channels than that of MUA and ESA. Compared to SUA and MUA, ESA needed a higher number of LFP channels to reach the maximum inference accuracy (shown by the red star markers in Figure 5d-e). This may indicate that ESA has broader spatial coverage and more information than SUA and MUA.

### Performance evaluation on different subject and task

Lastly, we next sought to determine whether the trend in the above results was associated with any particular subject or task. We performed the same inference procedure using a different subject performing a different task (see Dataset II in Methods). Analysis of LFP feature informativeness revealed the same trend where LMP yielded the highest inference accuracy, whereas alpha yielded the lowest inference accuracy. However, there was a minor difference in the order of LFP feature informativeness in which delta feature showed better predictive performance than gamma. The order of LFP feature informativeness (from highest to lowest) was as follows: LMP > delta > gamma > beta > theta > alpha (Figure 6a). The statistical tests among pairs of LFP features also showed that there were statistically significant differences among the features (Figure 6b). The relative performance of LMP varied from 1.28 to 10.53 times better than that of other features (Figure 6c). Using LMP as the feature, we plotted the inference accuracy of each channel of SUA, MUA, and ESA into 10-by-10 heatmap grids (see Figure 6d-f). The inference accuracy of ESA (average CC 0.702 ± 0.014) was also found to be statistically significant higher than SUA (0.552 ± 0.020) and MUA (0.525 ± 0.021) as illustrated in Figure 6g. Relative to SUA and MUA, ESA yielded on average 1.28 and 1.35 times better inference accuracy. Similar to the results obtained from the dataset I, the inference accuracy of SUA reached a plateau in less number of LFP channels than MUA and ESA (Figure 6h). In good agreement with the results from the dataset I, the LFP from the same channel had high but not necessarily the highest importance score for the inference of spiking activity (Figure 6i-k). Moreover, for the same channel, the importance score of ESA, on average, was also higher than that of SUA and MUA.

**Figure 6.**
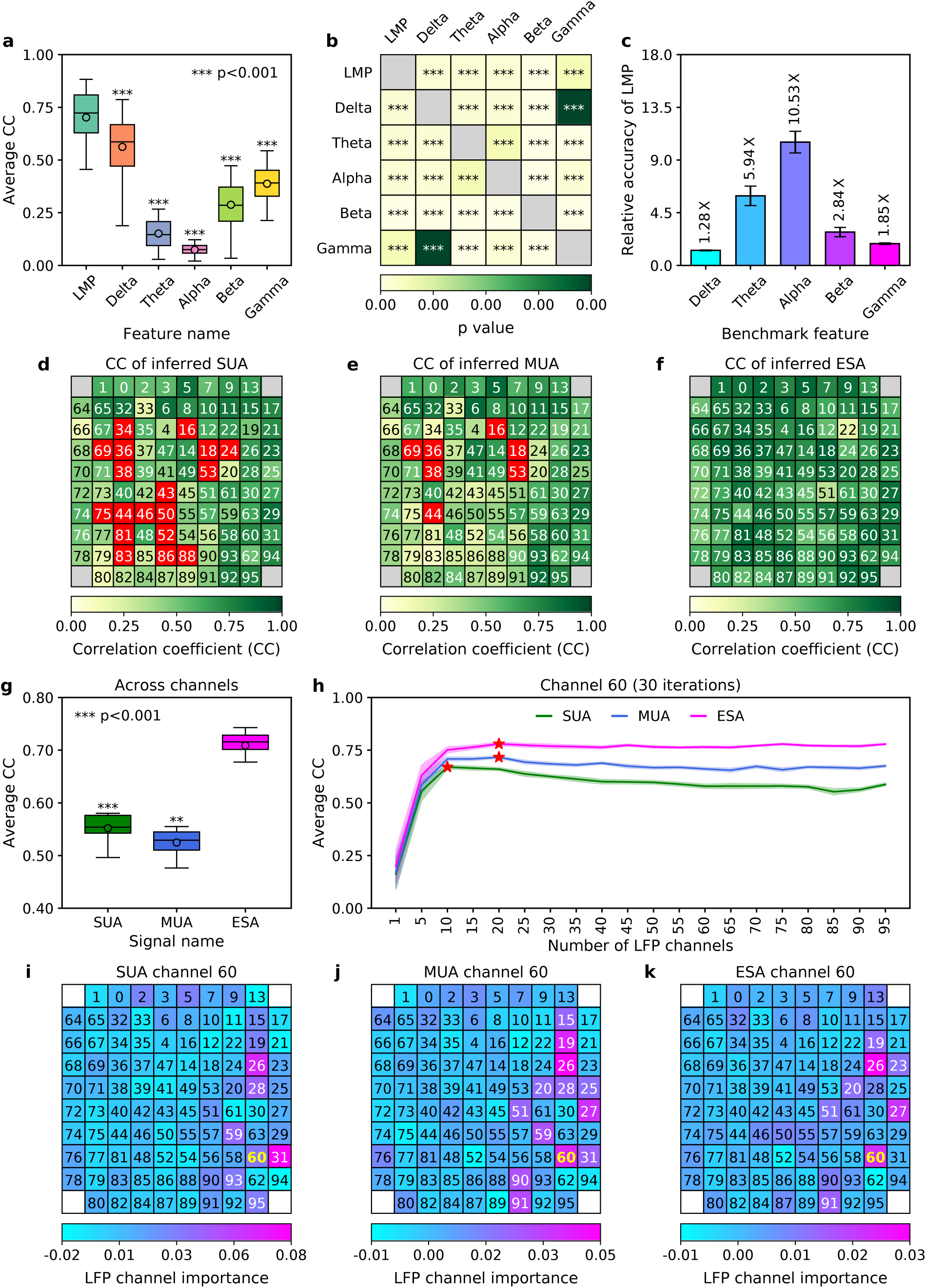
Performance evaluation on different subject and task (recording session I140703-001) (**a**) Boxplot comparison of ESA inference (average CC) across LFP features. Asterisks indicate LFP features whose inference performance differed significantly from that of LMP (*** *p*<0.001). (**b**) Heatmap of statistical significance matrix among LFP features. Grey boxes in the main diagonal does not have *p*-value (excluded from statistical test calculation). (**c**) Relative inference accuracy of LMP with respect to other LFP features. Larger value indicates better relative performance. Black error bars denote 95% confidence intervals. (**d**-**f**) Heatmap of CC from 96-channel of SUA, MUA, and ESA, respectively mapped onto a 10 ×10 grid spatially corresponding to Utah electrode array configuration. Black/white numbers inside the grids denote each of the 96-channel numbers. Grey boxes on the grid corners represent unused (unconnected) electrodes. Red boxes indicate channels that were excluded from the inference due to too few number of recorded spikes (≤ 0.5 Hz). (**g**) Boxplot comparison of average CC across channels among SUA, MUA, and ESA. Asterisks indicate the statistical significance of other spiking activities being compared to ESA (*** *p*<0.001). (**h**) The effect of increasing number of LFP channels for the inference accuracy (average CC) of spiking activities in channel 60. The shaded areas represent 95% confidence intervals (*N* = 30). The red star markers indicate the number of LFP channels that corresponds to highest average CC score for each spiking activity. (**i**-**k**) Heatmap of LFP channel importance score for the inference of SUA, MUA, and ESA in channel 60 mapped onto a 10 ×10 grid spatially corresponding to Utah electrode array configuration. Black/white numbers inside the grids denote each of the 96 LFP channel numbers. Bold yellow number (i.e. 60) indicates the channel of spiking activities being inferred. White boxes on the grid corners represent unused (unconnected) electrodes.

## Discussion

The present study investigates the relationship between local field potential (LFP) and entire spiking activity (ESA) by asking whether we can infer ESA solely from LFPs. In doing so, we firstly examined which feature within LFPs contains the highest information (i.e. most predictive) about ESA. Our experimental results revealed that local motor potential (LMP), the smoothed time-domain amplitude of LFP, emerged as the most predictive feature, followed by power feature either from delta or gamma frequency bands. On the other hand, the intermediate frequency bands of LFPs carried only a little information about ESA. A considerable number of previous studies have reported that power modulation within low frequency (< 10 Hz) and high-frequency (*>* 30 Hz) of LFPs were highly informative, whereas the intermediate frequency bands were little informative about spiking activity (SUA or MUA)^24, 30–33^. It is believed that the low-frequency band LFPs are associated with motor evoked potentials that usually exhibit high correlation with the spiking of neuronal population. On the other hand, the high-frequency band LFPs are thought to reflect synchronised spiking activity arising from local neuronal population^30^. Our study confirms and extends the previous findings by showing that apart from power within low- and high-frequency LFPs, LMP also contained substantial information about spiking activity. LMP even yielded highest inference accuracy in both CC and RMSE metrics across recording sessions and behavioural tasks. Even though LMP has been frequently used as a feature for decoding behavioural tasks^34–36^, the use of LMP for inferring spiking activity has not been investigated. A rather similar feature to LMP is low-frequency LFP (lf-LFP) which is obtained by low-pass filtering the broadband LFPs^26, 28^. The frequency band of lf-LFP is more clearly separated than that of LMP as low-pass filter results in better roll-off and stopband attenuation. On the other hand, LMP is obtained by a moving average filter which is simple, fast and yields less random white noise while maintaining well the smoothing of LFP signals^37^.

Given the highly informativeness of LMP, one may ask what the biophysical origin of LMP is. Unfortunately, the answer to this question remains to be established. It has been speculated that LMP is related to firing rate modulation of neuronal population near the recording electrode^38^ or distant neuronal population activity that is connected to the recording site^24, 39^. It is also possible that LMP reflects evoked LFPs in the motor cortex in response to movement-related tasks^30, 38^. A prior study suggested that LMP plays an important role not only in the execution but also in the preparation of movement (e.g. anticipation)^40^. This may explain the better relative inference accuracy of LMP with respect to other features from dataset II (movement with preparatory delay) than from dataset I (movement without delay interval).

Next, using the LMP feature and LSTM model with optimised hyperparameters, we evaluated the inference accuracy of ESA and compared its performance with that of SUA and MUA. Results across recording sessions, subjects, and tasks showed that the inference accuracy of ESA was consistently higher than that of SUA and MUA. Previous studies have only conducted the inference of SUA and MUA from LFPs^26, 28, 30–32^. Therefore, to the best of our knowledge, our study is the first that evaluates the inference of ESA from LFPs and systematically compares its performance to that of SUA and MUA. We argue that the high inference accuracy of ESA is attributed to richer spiking information contained within ESA compared to SUA and MUA. Both SUA and MUA are obtained by using a threshold-based technique which is prone to false-positive or false-negative spike detection. If we set the threshold value too high, we could miss true but lower amplitude spikes; on the contrary, if we set the threshold too low, we may detect some background noises as spikes. In contrast to SUA and MUA, ESA does not use thresholding, instead, it employs full-rectification and low pass filter which is rather insensitive to random high-frequency noise and is able to preserve full information of spiking activity^14^. Moreover, thresholding favours large (pyramidal) neurons with high amplitude spikes and other neurons very close to the electrode tips^13, 14^. The small neurons (with low amplitude spikes) or slightly farther neurons may not be detected. In contrast to that, ESA integrates the contribution from all neurons in the vicinity of the recording electrode tip.

To gain more insights into the relationship between LFP and spiking activity, we measured and ranked the LFP channel importance/contribution for the inference of SUA, MUA, and ESA at particular channels using feature importance permutation. The results revealed that the LFP importance score in the same channel as the inferred spiking activity on average had high (not necessarily highest) importance score, indicating its rich information about the spiking activity. This is in good agreement with the previous findings^24, 32^. Furthermore, within the same channel, the LFP importance score for ESA was on average higher than SUA and MUA. This may imply that ESA contains additional spiking information that is not detected by thresholding procedure in the cases of SUA and MUA. Lastly, we investigated the effect of a different number of LFP channels on the inference accuracy of spiking activity. We found that SUA and MUA achieved plateau and peak accuracy in less number of channels compared to ESA. To attain the peak inference accuracy, a larger number of LFP channels was needed for the inference of ESA. This could be an indication of broader spatial coverage of ESA (encompassing smaller and farther neurons) compared to SUA and MUA. These results may suggest that LFP represents the total input giving rise to spiking output within broader spatial area (observed through ESA) than previously thought (SUA or MUA).

In this study, we inferred spiking activity from LFPs recorded from the same electrode. This could confound our analysis since spike waveforms from nearby neurons may contaminate the LFPs. To remove this possible contamination, we used a filter with lower frequency cut-off (100 Hz) than the typical frequency cut-off (300 Hz) to obtain LFPs from the raw neural data. Although slow modulation of spiking activity as low as 10 Hz can contribute to the LFPs^30^, previous studies suggested that its contribution is negligible compared to the other sources such as synaptic activity and membrane potential oscillations^22, 41^.

In summary, we have shown that LFPs, in the form of LMPs, carry substantial information about spiking activity, particularly ESA. Since spiking activity has been widely used as an input signal for BMI, our finding corroborates the increasingly accumulating evidence that LFPs can be used as an alternative input signal. This is especially relevant for chronic recordings where the spike signals have been found to be unstable or even degrading over long periods of time. In this case, LFP-based BMIs can be implemented using biomimetic or biofeedback approach^12, 28, 42, 43^. Another finding from this study is that ESA appears to contain richer information about and broader spatial coverage of spiking activity within an area compared to SUA and MUA. This suggests that ESA could be potentially used in neuroscience research as an alternative neuronal population activity measure for analysing neural responses to various stimuli or behavioural tasks.

## Methods

### Neural recording and behavioural task

Electrophysiological recordings used in this study were obtained from two public neural datasets^44, 45^ herein referred to as dataset I and dataset II. These datasets were recorded from motor cortex area of two adult male Rhesus macaque monkeys (*Macaca mulatta*) while performing predefined tasks with 96-channel silicon-based intracortical microelectrode (Utah) array. The data acquisition and behavioural task associated with each dataset are briefly described below.

#### Dataset I

This dataset was recorded from a monkey (named as monkey I) while performing a point-to-point task, that is, to reach randomly drawn circular targets which were uniformly distributed around an 8-by-8 square grid. A sequence of new random targets was presented immediately and continuously after target acquisition without an inter-trial interval. The recordings were amplified and filtered with a 4th-order 7.5 kHz low-pass filter with a roll-off of 24 dB per octave. After that, the recordings were digitised with 16-bit resolution at 24.4 kHz sampling rate. These digitised recordings are hereinafter called as raw neural signals. More detailed description on the experimental setup is described elsewhere^46^. In our experiments and analyses, we used a total of 26 recording sessions spanning 7.3 months between the first (I20160627_01) and last (I20170131_02) sessions. The recording duration varies from 6 to 13.6 minutes with an average of 8.88 ± 1.96 minutes.

#### Dataset II

This dataset was recorded from a monkey (named as monkey N) while performing an instructed delayed reach-to-grasp task. The monkey had to grasp an object with one of grip types (side grip or precision grip) and displace it with either a high or low pulling force. In each trial, the monkey had to perform one of four possible trial types (from combination of two grip types and two force types) randomly drawn from an equiprobable distribution. Before initiating the movement, the monkey had to wait for 1000 ms (preparatory delay). The trial was successful if the monkey could reach, grasp, pull and hold the object for 500 ms within the position window. The recordings were amplified and filtered with a 1st-order 0.3 Hz high-pass filter and a 3rd-order 7.5 kHz Butterworth low-pass filter. The recordings were then digitised with 16-bit resolution at 30 kHz sampling rate, which hereinafter called as raw neural signals. This dataset contains only one recording session (identified as I140703-001) with a duration of 16.43 minutes. A comprehensive description on the experimental setup and behavioural task along with the corresponding metadata is provided elsewhere^47^.

### Signal processing and feature extraction

Signal processing steps from the raw neural signals to obtain LFP, ESA, MUA, and SUA along with their associated features are illustrated in Figure 1a–d. We briefly describe the signal processing and feature extraction steps as follows.

#### Local field potential (LFP)

LFP was obtained by low-pass filtering the raw neural signal with a 4th-order Butterworth filter at 100 Hz and then downsampling it to 1 kHz. 100 Hz cut-off frequency was selected to eliminate possible contamination from multiunit spiking activity within higher LFP band (100 −300 Hz). The filtering was performed on the forward and backward directions to avoid any phase shift. We extracted six different LFP features consisting of average amplitude feature called local motor potential (LMP) and average spectral power features in five different frequency bands: delta (0.5 −4 Hz), theta (4 −8 Hz), alpha (8 −12 Hz), beta (12 −30 Hz) and gamma (30 −100 Hz). The LMP was calculated using time-domain moving average filter with 256 ms rectangular window. The spectral power in each band was computed by applying short-time Fourier transform (STFT) with a 256 ms Hanning window to narrow band LFP signal (i.e. band-pass filtered LFP with 3rd-order Butterworth filter). The LMP and spectral power features were extracted through an overlapping fashion to yield a sample every 4 ms.

#### Entire spiking activity (ESA)

ESA was obtained by first digitally re-referencing the raw neural signals with common average reference (CAR) and high-pass filtering with 1st-order Butterworth filter at 300 Hz. The filtered signals were then full-wave rectified, low-pass filtered with 1st-order Butterworth filter at 12 Hz and downsampled to 1 kHz. All the filtering processes were performed in both forward and backward directions. We extracted one ESA feature using time-domain moving average filter with a rectangular window of 256 ms width and 252 ms overlap (similar to that of LMP feature).

#### Multiunit activity (MUA)

Spike waveforms were extracted by first band-pass filtering the raw neural signals with Butterworth filter (4th-order, from 500 Hz to 5kHz for dataset I; 2nd-order, from 250 Hz to 5 kHz for dataset II) and then storing snippets of the filtered signals (48 or 64 samples for dataset I; 38 samples for dataset II) that crossed a threshold value. These extracted spike waveforms would be used later for classifying spikes into distinct putative single units (i.e. spike sorting). In each channel, MUA was defined as all the detected spikes and represented by their spike times. Details on the spike detection for each dataset has been described in other studies^46, 47^. We extracted spike rate feature by convolving each MUA’s spike train with a 256 ms Gaussian window (standard deviation of 110 ms). This feature extraction was performed in an overlapping fashion to obtain feature sample every 4 ms. Only MUAs with spike rates exceeding 0.5 Hz were included for our experiments and analyses.

#### Single-unit activity (SUA)

SUA was obtained by aligning the extracted spike waveforms, reducing the dimensionality to a few principal components, and then sorting them into single units via certain algorithms (operator defined templates for dataset I; K-Means and Valley Seeking for dataset II). Details on how spike sorting was performed for each dataset can be found in other studies^46, 47^. We computed spike rate from SUA using the same method as for MUA. In this study, we only used one SUA per channel. If a channel had more than one SUAs, one representative of SUAs with highest spike rate was selected. In addition, only SUAs with spike rates above 0.5 Hz were included.

### Long short-term memory (LSTM)

Long short-term memory (LSTM), proposed by Hochreiter and Schmidhuber in 1997^48^, is one of the most popular deep learning architectures and has achieved state-of-the-art performance in a wide range of machine learning problems, especially those dealing with time-series data^49^. It addresses the problem of vanishing or exploding gradient commonly found in traditional recurrent neural networks (RNNs). LSTMs can effectively learn long-term temporal dependencies via a memory cell that maintains its state overtime and gating mechanism that controls the flow of information into and out of the memory cell. In this study, we employed a commonly used variant of LSTM architectures which consists of three gates (forget, input, output) and a single memory cell. The states of LSTM components at timestep *t* are formulated as: 

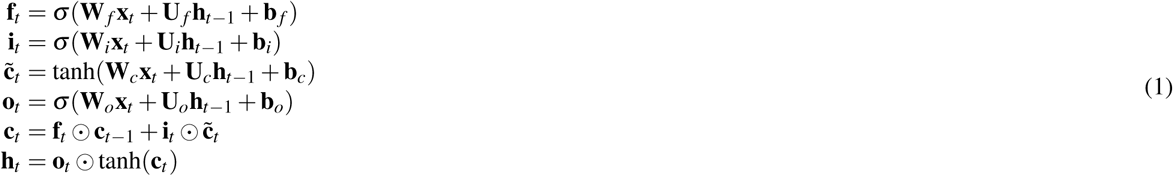

where **x, h, f, i, o, c** consecutively represent the input, output, forget gate, input gate, output gate, and memory cell. *σ* and tanh denote the logistic sigmoid and hyperbolic tangent activation functions, respectively. The operator ⊙ denotes the element-wise multiplication. Parameters that are learnt during training include input and recurrent weights (**W, U**) and bias vectors (**b**). We used 1-layer LSTM with number of timesteps (i.e. sequence length) empirically set to 2. Other hyperparameters including number of units, number of epochs, batch size, dropout rate and learning rate were determined through hyperparameter optimisation (described in the following subsection). The last timestep from the LSTM output was connected to a fully connected layer to obtain the final output. The LSTM model was implemented using Pytorch (v1.3.1) deep learning framework^50^.

### LSTM model training and optimisation

We divided each dataset into 10 non-overlapping contiguous blocks of equal size which were then categorised into three sets: training set (8 concatenated blocks), validation set (1 block) and testing set (1 block). The training set was used for training the model, the validation set was used for optimising the hyperparameters, whereas the testing set was used for the final performance evaluation. The schematic of the LSTM model training, optimisation and evaluation is illustrated in Figure 1e,f. We standardised (i.e. z-transformed) the LFP and spike features to have zero mean and unit standard deviation. The model was trained using RMSprop optimiser and mean squared error (MSE) loss function. We used Bayesian optimisation library called Hyperopt^51^ to optimise the LSTM hyperparameter values from predefined hyperparameter search spaces shown in Table 1. We optimised the hyperparameters for inferring ESA, MUA, and SUA signals separately using the same procedure. In the case of ESA, the hyperparameter optimisation was performed on each of six different LFP features. In the cases of MUA and SUA, the hyperparameter optimisation was only performed on the LMP feature of LFP. The optimised hyperparameter values for each case are provided in Table 1.

**Table 1.** Hyperparameter setting of LSTM models for inferring different types of spike signals. For inferring ESA signal, hyperparameter values were optimised separately on each of six LFP features. For inferring MUA and SUA signals, hyperparameter values were optimised on the LMP feature of LFP.

| Hyperparameter | Value range | Optimised value for different signal types |  |  |  |  |  |  |  |
| --- | --- | --- | --- | --- | --- | --- | --- | --- | --- |
|  |  | ESA |  |  |  |  |  | MUA | SUA |
|  |  | LMP | Delta | Theta | Alpha | Beta | Gamma |  |  |
| Number of units | $\{5, 10, \dots, 25\}$ | 25 | 15 | 5 | 5 | 10 | 25 | 25 | 25 |
| Number of epochs | $\{2, 3, \dots, 7\}$ | 2 | 3 | 5 | 4 | 4 | 2 | 2 | 3 |
| Batch size | $\{32, 64, 96\}$ | 32 | 64 | 96 | 64 | 96 | 96 | 64 | 64 |
| Dropout rate | $\{0, 0.1, \dots, 0.4\}$ | 0.2 | 0.4 | 0.4 | 0.4 | 0.2 | 0.4 | 0.2 | 0.3 |
| Learning rate | $\{5, 10, \dots, 50\} \times 10^{-4}$ | 0.002 | 0.001 | 0.0005 | 0.0005 | 0.0005 | 0.0005 | 0.001 | 0.001 |

### LFP channel importance/contribution

The importance/contribution of each LFP channel on inference performance was measured using a method introduced by Breiman in 2001^52^ known as permutation importance. Specifically, we used a model-agnostic version of the feature importance proposed by Fisher *et al.* which they call model reliance^53^. It measures the importance of a channel (i.e. variable) by computing the difference of model’s inference accuracy after temporally permuting the data in that channel. This permutation breaks the temporal relationship between the data in that channel and output variable being inferred, which increases the model’s accuracy difference. The larger the increase in the model’s accuracy difference after channel permutation, the more important is that channel for the model’s inference.

The procedure to calculate the channel importance is as follows: *first*, compute the accuracy of the original trained model (without permutation) based on the testing set and record it as the baseline accuracy; *second*, for each channel, permute the data in that channel, compute and record the accuracy based on the permuted data; *lastly*, compute the channel importance as the difference between the baseline accuracy and the accuracy based on the permuted data. We implemented this procedure using a Python library called MLxtend^54^.

### Effect of number of LFP channels on inference accuracy

To examine the effect of number of LFP channels on the inference accuracy, we selected randomly *n* distinct LFP channels where *n* = {1, 5, 10, 15, *…*, 90, 95} from a total of 96 channels. We then used these *n* LFP channels to train LSTM model evaluate its performance in inferring SUA, MUA, and ESA. This procedure was repeated for 30 iterations to obtain the average performance along with its confidence interval. The inference accuracy of SUA, MUA, and ESA from the same channel number were compared and analysed.

### Performance evaluation and metrics

The performance evaluation of LFP to spiking activity (ESA/MUA/SUA) inference was conducted using the optimised LSTM model on the testing set with Pearson’s correlation coefficients (CC) metric. CC measures the linear correlation between the actual and inferred spiking activity and has been used in several prior studies^26, 28, 33, 55, 56^. The formula for computing CC is as follows: 

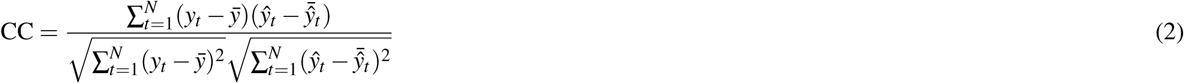

where *y*_*t*_ and *ŷ*_*i*_ represents actual and inferred spiking activity at time step *t*, respectively and *N* is the total number of samples. In addition, we also evaluated the inference performance using another metric called root mean square error (RMSE). It is a measure of the average magnitude of the inference error and has also been previously used in other studies^26, 33^. RMSE is formally defined as follows: 

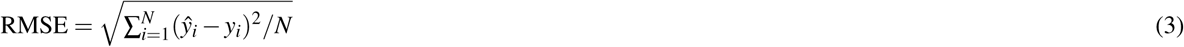

For each session, the mean and confidence interval of the inference performance were evaluated on 10 different blocks within the testing set. To test for significant effects between a pair of different LFP features or different spiking activity, we used a paired two-tailed *t*-test whenever the difference between the pairs follows normal distribution, otherwise we used Wilcoxon signed-rank test. The significance level (*α*) was set to 0.05.

When we visualised the result with boxplot, the horisontal line and circle mark inside each box represent the median and mean, respectively. The colored solid box represents interquartile range (from 25th to 75th percentiles). The whisker extends 1.5 times the interquartile range. All the analysis were conducted in Python (v3.6.9).

## Data Availability

Data are available from Zenodo at https://zenodo.org/record/583331 and from the German neuroinformatics node’s data infrastructure (GIN) at https://gin.g-node.org/INT/multielectrode_grasp.

## Acknowledgements

This work was supported by the UK Engineering and Physical Sciences Research Council (grant number EP/M020975/1) and Indonesia Endowment Fund for Education (LPDP) graduate scholarship program (grant number PRJ-123/LPDP/2016).

## Author contributions

N.A., T.G.C. and C.S.B. conceived the study, N.A. developed the methodology, conducted the experiments, analysed and visualised the data, N.A., T.G.C and C.S.B interpreted the results, N.A. wrote the original draft. All authors reviewed and edited the manuscript.

